# *demografr*: A toolkit for simulation-based inference in population genetics

**DOI:** 10.64898/2025.12.18.694482

**Authors:** Martin Petr, Isabel M. Pötzsch, Fernando Racimo

## Abstract

Simulation-based inference methods such as Approximate Bayesian Computation (ABC) are a popular class of techniques in evolutionary biology and population genetics. These methods are particularly useful for fitting complex models, as they can bypass the need to compute an exact likelihood function, instead relying on comparisons between observed and simulated summary statistics. However, numerous technical and practical hurdles often hinder the application of these methods to new modeling problems. They require the integration of disparate scripts and software for simulation, computing summary statistics, and inference into pipelines which have to be tailor-made for each specific research project. Moreover, computational costs of population genetic simulations can make inference laborious due to costly file format conversions throughout the entire procedure. Here we present a new R package called *demografr* (github.com/bodkan/demografr), which aims to alleviate most issues with traditional simulation-based inference workflows while facilitating their application to novel and complex modeling problems. First, *demografr* leverages the R simulation toolkit *slendr* for interactive, user-friendly encoding of complex demographic models. Second, *demografr* allows the user to set up parameter distributions and run simulations from them automatically, with built-in support for parallelization. For this purpose, automated inference routines for ABC or grid-based parameter exploration are provided, with individual components available for developing other simulation-based workflows in the future. Third, *demografr* allows a wide range of summary statistics to be efficiently computed directly in R, without any need for file conversion. In addition to the default *slendr* simulation engine, *demografr* also supports user-defined *msprime* and SLiM simulation scripts as well as entirely customizable summary statistic functions. We illustrate the features of *demografr* on examples which would traditionally require complex workflows with hundreds of lines of code: estimating parameters of a demographic model via coalescent simulations using ABC, exploring the influence of a set of parameters on summary statistics of interest using parameter grid search, and showcasing the ability to integrate fully customized simulation code written in pure Python or SLiM.

## Introduction

Simulation-based inference methods have proven to be invaluable for inference across the sciences [1] and, in particular, in population genetics and evolutionary biology [2]. Broadly speaking, they are based on simulating genetic data from models sampled from given parameter distributions, and then comparing these simulations to observed data, retaining the values of parameters which are most consistent with the observations. Parameters which are often estimated in these types of frameworks include effective population sizes (*N_e_*), migration rates, or divergence times. The flexibility of simulation-based methods allows exploring a potentially much broader range of demographic scenarios than standard likelihood methods [3,4], both in terms of the number of populations and their topological relationships that can be evaluated, as well as the number of individual demographic events in their evolutionary histories. This flexibility makes simulation-based methods well-suited for situations in which explicit likelihood functions may be challenging to derive and, as such, present a powerful and versatile alternative for model fitting. Among the most popular methods for simulation-based inference are Approximate Bayesian Computation (ABC) [2], the coalescent-based inference method implemented in *fastsimcoal2* [5], or even simple grid-search parameter approaches [6,7].

Despite their utility and wide applicability—especially with the increasing complexity of models driven by the growing amount of ancient and modern genomic data available across species [8–10]—simulation-based methods remain relatively underutilized for several practical reasons.

The first issue involves model specification. Simulation-based inference requires a researcher to program custom-tailored simulation code which captures the essential features of the model in question (Figure 1A). Although great progress has been made in the development of powerful, fully-programmable libraries and frameworks for programming models in population genetics [11,12], developing simulation code remains a significant hurdle for researchers who are not comfortable with coding. Unfortunately, the amount of programming needed for all but the most trivial models is quite significant (easily reaching several hundreds of lines of code), which increases the occurrence of software errors even for computationally-savvy researchers [13]. Worse still, every new modeling study utilizing simulations requires researchers to reimplement simulation scripts effectively from scratch, tailored for species and modeling questions at hand. This duplication of effort dramatically increases the amount of programming work needed across different projects and, consequently, the occurrence of software errors across the entire field. This is frustrating because, arguably, many demographic models in population genetics are composed from a limited number of well-defined events such as population splits, changes in population sizes, and gene flows and thus are, apart from the tree topology of the base demographic history, effectively a variation on the same theme [8,9]. A recent unified effort for an increase in reproducibility has pushed for a more standardized method of encoding published models [14,15], however, an easy-to-use method for encoding parametrized models (that is, models with unknown parameters values which have to be inferred from data) is still missing. This issue is even more pressing for spatially-explicit models, which can involve much larger parameter spaces, are significantly more complex and, as a result, even more challenging to work with [16].

**Figure 1.**
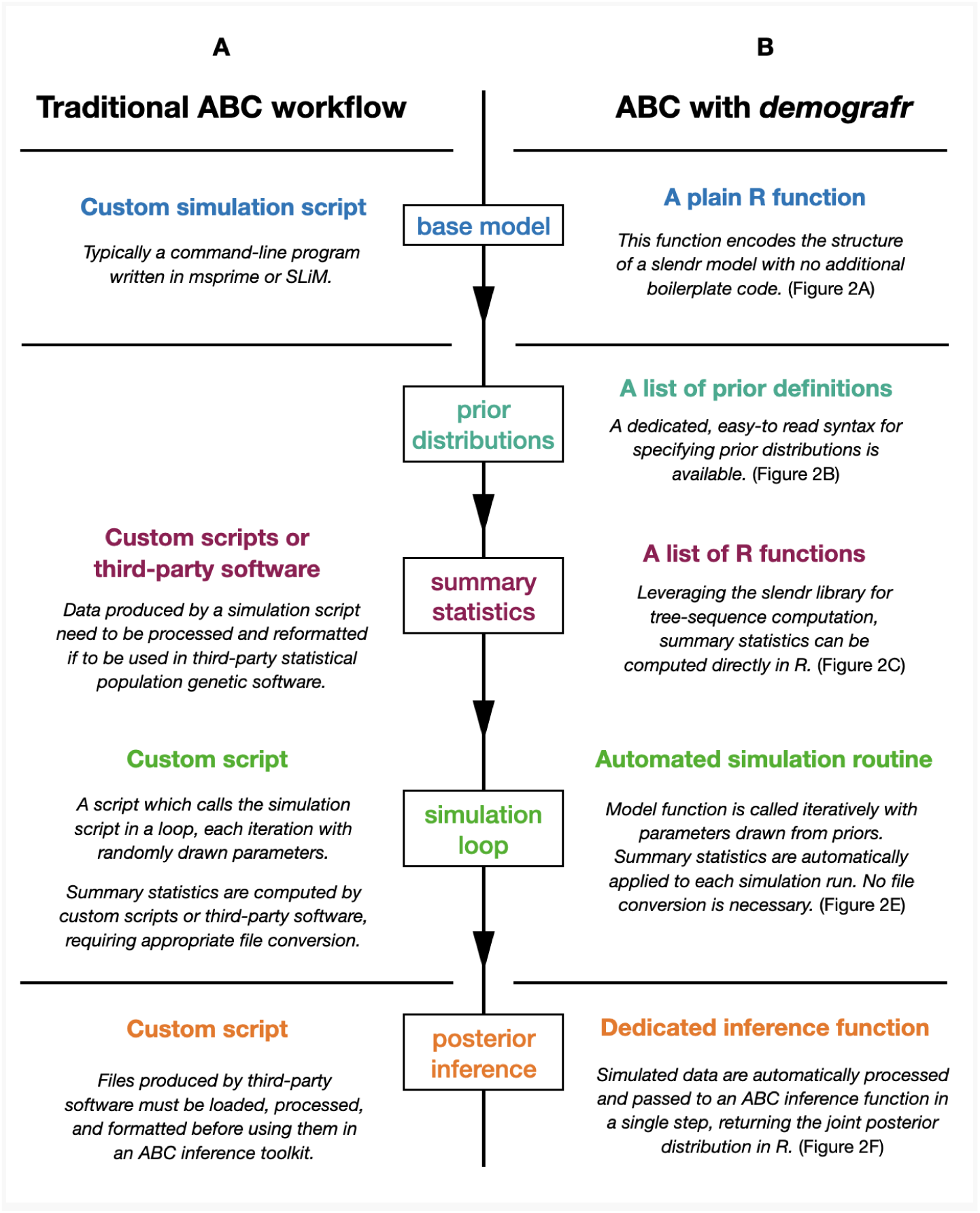
A schematic diagram comparing the typical steps (and the respective software components thereof) of a traditional ABC inference workflow (A) and a corresponding version of the same workflow using *demografr* (B).

Second, after simulation code has been written and executed, summary statistics of interest need to be computed from the simulations (Figure 1A). In the most straightforward case, for each simulation replicate of a particular parametrization of a model, result in the form of a VCF file [17] or a customized genotype table data file is typically saved to disk, and potentially further converted to other relevant formats such as EIGENSTRAT [18] or PLINK [19], depending on the software used to compute the summary statistics of interest. Because disk access is typically much slower compared to the simulations themselves, this can become potentially prohibitive, in particular for computationally heavy methods such as ABC which can routinely require millions of simulations [20]. Creative approaches of a low-level streaming of simulated genotypes directly to memory have been used in the past for the most extremely intensive simulations [21] but these require custom, low-level modification of simulation software, and are thus not generalizable to all possible scenarios and all possible summary statistics. Additionally, aside from slow disk serialization of simulation results, computing population genetic statistics from them has traditionally required building customized pipelines for integration into external software such as ADMIXTOOLS, PLINK, or ANGSD [18,19,22]. This further compounds on the above-mentioned issues of reproducibility and the propensity for errors to occur and proliferate with each new project [13].

Third, once custom code for simulating data from a model and computing summary statistics is written, a pipeline which will integrate both of these components together to form an overall inference workflow needs to be developed (Figure 1A). For a traditional ABC analysis, this typically requires yet another custom-tailored script which will sample parameters from prior distributions in a loop, execute the simulation with those parameters, format the simulated data appropriately, and compute an array of summary statistics across sampled parameters. Individual simulated summary statistics are then gathered and used in ABC inference to estimate the posterior distribution of parameters, either reformatted in data format expected by dedicated inference engines such as the popular R package *abc* [23] or via simpler ABC-like acceptance-rejection algorithms [24]. Dedicated software pipelines avoiding some of these issues have been developed [25], however, these do not allow the use of modern simulation tools necessary for the genome-scale data available today [11,26,27]. Other simulation-based techniques, including a simple parameter sweep across a grid of predefined values, largely follow a similar workflow requiring extensive custom scripting [6].

Thus, although powerful and flexible, typical simulation-based inference workflows require a significant amount of programming effort across various (yet otherwise very similar) modeling studies, which leads to a frustrating degree of code reimplementation (Figure 1A). Consequently, given the increasing complexity of models under study, this need for repeated code reimplementation significantly increases the opportunity for software errors and hinders reproducibility [28]. Here, we present an inference toolkit implemented in the R package *demografr*, which addresses all these problems in several ways. First, by utilizing the functionality of the simulation framework *slendr* [29] for encoding demographic models using its straightforward domain-specific syntax, it reduces the amount of coding necessary to program demographic models down to writing a single plain R function without any additional boilerplate code. Second, because *demografr* models produce simulation results as a tree-sequence object in the *tskit* format, a wide array of summary statistics can be computed trivially using *slendr*’s interface to the *tskit* module [26]. Third, *demografr* provides several routines for automatic, parallelizable execution of various demographic inference methods, replacing the need for writing custom code. It does so by calling a single function which interfaces with the popular R package *abc*, allowing users to leverage its functionality for model selection and diagnostics naturally, without any additional work. Finally, by providing individual building blocks for any general simulation-based inference workflow (i.e. defining a model, parameter distributions, summary statistics, and simulating data), routines for other inference paradigms can be developed to expand the range of *demografr*’s inference functionality in the future.

### Installation

A development version of *demografr* can be installed from GitHub using the R package *devtools* as `devtools::install_github(“bodkan/demografr”).` For convenience, a Docker container which includes all software dependencies and features a browser-based installation of RStudio (thus providing a complete self-contained research environment) is available on DockerHub as a container image `bodkan/demografr.`

### Design

In this section we describe the individual components of a typical *demografr* inference workflow (Figure 1B), explain how each step integrates with the *slendr* population genetic simulation toolkit, and showcase how to extend *demografr* pipelines to user-defined simulation scripts.

### Model specification

The *demografr* toolkit embraces the R package *slendr* as the default means to specify the structure of a demographic model (Figure 2A). Specifically, a model is encoded as a simple R function whose arguments represent parameters to be inferred. Later, during the simulation stage, values of these parameters will be drawn from prior distributions specified using *demografr*’s intuitive syntax (Figure 2B). Inside the model function, demographic components of *slendr* (such as `population()`, `resize()`, and `gene_flow()` functions) can be used to compose practically any kind of tree topology of traditional demographic models. Similarly, custom sampling schemes—i.e., scheduling “ancient DNA” sampling events or recording custom-defined subsets of simulated populations in general—can be included through *slendr*’s own `schedule_sampling()` calls. Finally, we note that because *slendr* can utilize both *msprime* and SLiM as simulation engines, and does not principally distinguish between non-spatial and spatial models, *demografr* supports spatial models with no additional changes to the modeling interface described here. Thus, by utilizing *slendr* as a model-definition framework (and, as we show below, also its simulation engine), *demografr* models can be expressed by users in a concise and readable manner, while retaining all the flexible functionality for model inspection and visualization provided by the underlying *slendr* package.

**Figure 2.**
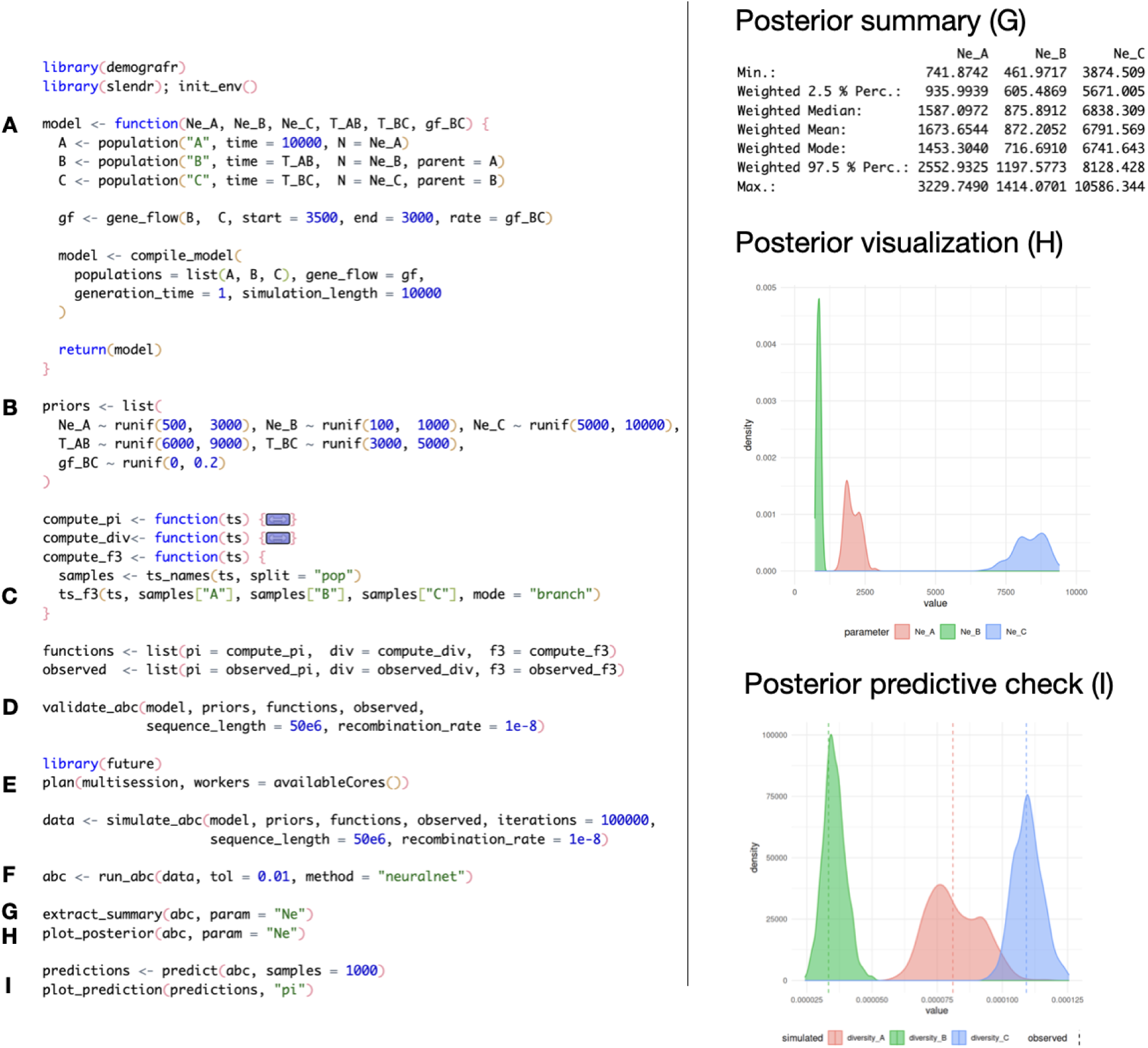
Basic workflow demonstrating individual steps (A-I) of a typical ABC inference R script implemented using the *demografr* package. Panels on the right show results produced by the commands G-I in the script on the left. Complete reproducible code for this example, as well as additional possibilities are shown in a corresponding vignette at https://bodkan.net/demografr.

### Custom models

Although *slendr* captures the most important components of traditional population genetic models as mentioned (i.e., creating populations, population splits, *N_e_* changes, gene flow events, assuming Wright-Fisher dynamics), its focus on user-friendliness and conciseness imposes certain restrictions on the vast array of features that SLiM or *msprime* could deploy on their own. For these situations, *demografr* allows users to provide custom-designed *msprime* scripts written in pure Python [11] or SLiM scripts written in Eidos [12]. The only requirement placed by *demografr* on such scripts is the ability to accept model parameters on the command-line (Figure 4J). This way, users have every feature of *msprime* and SLiM at their disposal, including those currently not supported by *slendr*. A demonstration of using *demografr* for inference using “pure” Python or SLiM scripts is shown in *Example 2*.

### Summary statistics

In addition to custom simulation code, simulation-based inference pipelines often require another layer of customized code for computing summary statistics (Figure 1A). At the very least, this generally involves file format conversion for computation in third-party software or scripts for computing summary statistics and, at a later stage, importing and reformatting the results for comparing empirical and simulated data. This can be quite time consuming both in terms of the time needed to serialize data to disk and because of the time required to write this data processing code itself.

Recently, an exciting new data structure called “succinct tree sequence” (hereafter “tree sequence” for short) has been introduced, representing an efficiency breakthrough in storage and computation for large-scale genomic data [26,27], organized around the software project *tskit* (www.tskit.dev). Because *demografr* leverages the *slendr* package for model definition and data simulation, the default outcome of each *demografr* simulation is a *tskit*-compatible tree sequence object. Furthermore, because *slendr* includes a built-in coalescent simulation engine written in *msprime* which operates entirely in memory, the tree-sequence data it produces is never saved to disk and summary statistics can be thus computed without any file manipulation. Finally, because *slendr* provides an R-idiomatic interface to the most common population genetic tree-sequence statistics implemented by *tskit* (site-frequency spectrum, *f*-statistics, F_ST_, nucleotide diversity, Tajima’s D, etc.) [29], users can specify which statistic should be applied to each simulation replicate and how directly in R, without having to do any additional work (Figure 2C).

For most common inference tasks, this approach completely eliminates the need to convert simulated data for computation in external software, allowing for both the tree-sequence data simulation and statistical computation to happen efficiently in a single R environment (Figure 1B). Furthermore, because *slendr*’s tree-sequence *tskit* functions produce results as standard R data frames, downstream inference involving comparisons to equivalent observed statistics is completely streamlined. In fact, owing to this feature, matching of simulated and observed statistics in the ABC context is done by *demografr* internally and automatically, without the user having to manually keep track of which column of which simulated statistic’s data frame corresponds to which part of the observed data, as is needed, for instance, by traditional pipelines built around the *abc* R package [23].

### Custom summary statistics

Similarly to the option to use non-*slendr* scripts as simulation “engines”, *demografr* allows computation of summary statistics using functions outside of the library of currently supported *tskit* functionality of the *slendr* package [29]. This can include *tskit* tree-sequence Python methods which do not yet have a corresponding R interface in *slendr*, but *demografr* also allows users to define summary statistics which do not operate on tree-sequence objects at all, such as simple tables of genotypes, VCF files, etc. Their conversion to a form which can be utilized by *demografr* is automatically performed by so-called “data-generating functions”, which are, again, specified as standard R functions, and are given as an argument to any of the simulation routines. This feature is demonstrated in *Example 3*.

### ABC inference

Once a model, parameter priors, and summary statistic functions are specified, users can proceed to data simulation and, finally, inference of posterior distributions (Figure 1B).

Following the definition of the model, priors, and summary functions objects in R, users can execute a given number of simulations with a single line of code, such as

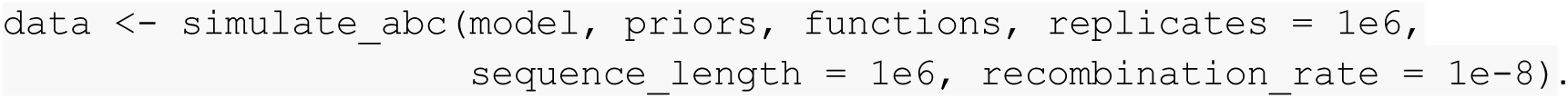

This example command would automatically execute one million simulations in a fully parallelized manner, each simulation replicate drawing parameter values from their `priors`, generating a given `model` with those parameters, simulating data (typically a tree sequence), and automatically computing all summary statistic `functions` on it. Thus, with this concise bit of R code, *demografr* encapsulates what are typically dozens of lines of customized code orchestrating simulations in a loop, abstracting the data-simulation process behind a single function call.

Finally, users can proceed to inferring posterior distribution parameters from the simulations. This can, again, be performed by executing a single command, such as

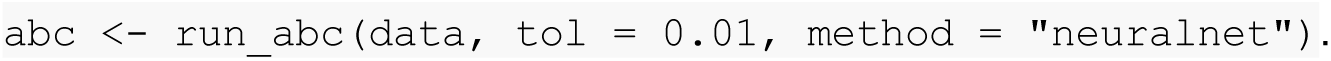

This function serves as a wrapper for the inference function `abc()` implemented by the *abc* R package [23], accepting all arguments normally available to this function: in this example, a tolerance of 1% and a choice of ABC inference algorithm based on neural networks [23]. Because the *demografr* `data` object encapsulates all information about every component established upstream from this step of the ABC workflow (i.e., the model used to generate data, priors, summary statistics, and all other simulation parameters), users do not need to do any additional work preparing and formatting data structures for inference, as is the case in traditional workflows using the *abc* package [23]. This is because all *demografr* wrappers internally extract all the necessary information, reformat it in the required way, and then call the “low-level” *abc* functions as needed, all behind the scenes. Although seemingly a technicality, less manual work, more automation, and a tighter link between code and data dramatically decrease the number of software bugs [28].

The resulting object `abc` represents the inferred posterior distribution which can be examined in several ways. First, because of *demografr*’s emphasis on backwards compatibility, all the visualization functions of the *abc* package can be utilized. Second, because the object encompasses all the necessary information about the pipeline that produced it, *demografr* also provides more user-friendly functions for extracting posterior summary statistics as standard R data frames (`extract_summary()`), extracting posterior samples (`extract_posterior()`), and plotting capabilities implemented using the *ggplot2* package (`plot_posterior()`) as shown in *Example 1* (Figure 2G-I).

### Parameter grid search

In addition to estimating the posterior distributions of parameters via ABC, *demografr* also supports simple grid-based parameter sweeps, which can be useful to investigate the behavior of a model under different interactions of its parameters [6]. Given a *slendr* model function and a list of tree-sequence summary statistic functions, just as described for the ABC case above, the user may perform such grid-based exploration across a desired range of simulation replicates by providing the combinations of parameters of interest as a standard R data frame to the function `simulate_grid`() as shown in *Example 2* (Figure 3). The results of this procedure are returned in another data frame, with the outcome of each simulated summary statistic for each replicate stored in a list-column format in intuitively named data-frame columns. This concise and readable representation allows straightforward extraction of simulated statistics across each parameter combination using tools for working with *tidy* data [30]. Again, the entire procedure is automated, parallelized, and the only components of the workflow that need to be provided are a *slendr* model function, a list of summary statistics, and parameter grid values to simulate across.

**Figure 3.**
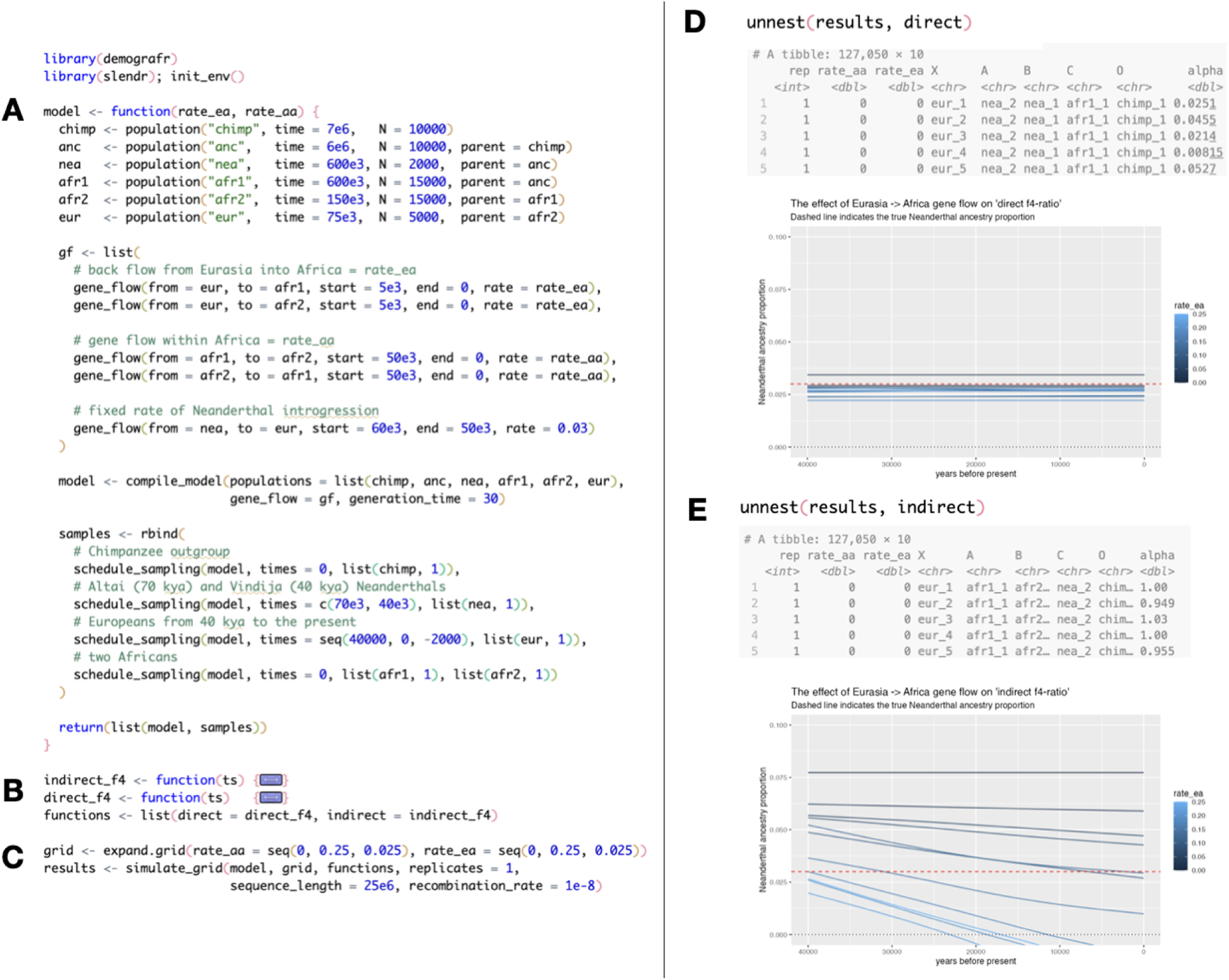
Basic workflow demonstrating individual steps (A-C) of a typical parameter grid search R script implemented using the *demografr* package. Panels on the right show the format of results produced by the simulation procedure in step C in the script on the left, as well as their immediate graphical representation. Complete reproducible code for this example, as well as additional possibilities are shown in a corresponding vignette at https://bodkan.net/demografr.

### Support for new inference routines

The `simulate_abc()` / `run_abc`() and `simulate_grid`() inference routines above have been showcased due to their popularity in population genetics, and for their immediate applicability to ongoing research workflows. However, a number of other statistical inference methods could be beneficial in certain settings, such as Bayesian parameter optimization [31], genetic algorithms [32], or random forests [33]. Although their application in the context of simulation-based inference has, to our knowledge, not been thoroughly explored, the building blocks for developing simulation-based workflows provided by *demografr* make it quite straightforward to implement analogous inference routines for each of them. For instance, a hypothetical simulation-based genetic algorithm inference routine would involve taking a model function, a prior parameter distribution, and summary statistics (building blocks already provided by *demografr*, Figure 1), integrating them into a standard R loop which would utilize the function `simulate_model()`, and applying a genetic algorithm procedure [34] to each simulation result. We envision that new inference routines of a similar kind could be developed in a community-developed library in the future.

### Parallelization

A characteristic feature of simulation-based inference techniques is their computational intensity. At the most extreme, an ABC workflow can easily require millions of simulation replicates [20], but all of these can be run entirely in parallel due to their mutual independence. To facilitate efficient parallelization with minimum amount of effort from the user, *demografr* leverages the R package *future* [35], which only requires the user to specify a “parallelization plan” (typically either multicore run on a single computer or distributed across multiple machines) at the beginning of their inference script. For instance, the R command `plan`(multisession, workers = 100) will instruct all downstream *demografr* simulation functions to distribute computation across 100 CPUs in a parallel manner automatically, without any additional work on the user’s part.

### Posterior predictive checks, model selection, diagnostics

A critical phase of any simulation-based inference workflow, particularly ABC, is model selection and diagnostics, such as posterior-predictive checks [36,37]. *demografr* makes these procedures straightforward, as the result of its ABC pipeline is ultimately a standard *abc* R object wrapped in a custom object class (Figure 2F). As such, *demografr* is fully interoperable and backward-compatible with every visualization, model selection, and model diagnostic functionality offered by the *abc* package. First, where convenient, *demografr* wraps such functionality in its own custom function (such as model_selection()) to eliminate the manual work required to be done by the user in traditional *abc* scripts, i.e. without having to keep track of vectors, matrices, or data frames of individual summary statistics, simulated and observed. All these components are already wrapped by *demografr* in a single all-encompassing result object, making it possible to streamline this process without any need for user intervention. That said, where necessary, *demografr* provides a helper function unpack(), which deconstructs this result object into its individual components, allowing for integration with any diagnostic function of the *abc* package directly while still effectively eliminating the chance for unintended software bugs. Finally, in addition to its compatibility with diagnostic functions of the *abc* package, *demografr* also provides an efficient, fully parallelizable method predict() for straightforward execution of posterior predictive checks which, paired with the function plot_prediction(), allows quick model iteration and troubleshooting.

## Examples

In this section, we provide several examples of workflows which we envision will be of most immediate interest to researchers in the broad field of evolutionary genetics. We note that the code for these examples was deliberately chosen to be as concise as possible, in order to emphasize the most important features of *demografr*, without exposing the reader to an unnecessary level of detail. However, more elaborate examples of these (and many other) use cases of *demografr* are discussed in tutorials available as standard R package vignettes on the project website (www.bodkan.net/demografr).

### Example 1: Basic ABC workflow

Figure 2 showcases the features of *demografr* on a textbook toy example of an ABC workflow, demonstrating the specification of a base model, definition of tree-sequence summary statistics, executing simulations, as well as an examination of posterior distributions and model diagnostics.

Briefly, model is a plain R function which internally uses the *slendr* interface to encode a simple three-population demographic model (Figure 2A). The model parameters to be inferred are always assumed to be the arguments of this function (Ne_A, Ne_B, , etc.). For brevity, the model does not feature population size changes or other, more complex, features such as spatial dimension.

The variable priors stores a list of “prior sampling expressions” using *demografr*’s domain specific syntax of <PARAMETER> ∼ <SAMPLING expression> (Figure 2B). Crucially, each name of a parameter specified in the prior must match exactly one of the model function arguments. In a later simulation stage, this allows *demografr* to internally sample parameters from their priors and call the model function in a fully automated way.

Because summary statistics operate on tree sequences by default, users provide simulated summary statistics to be computed on each simulation replicate as a list of R functions (Figure 2C), with the name of each element corresponding to one summary statistic, and expected to match a list of observed statistics (Figure 2C). Before proceeding with full-scale simulations, users have the opportunity to validate the entire pipeline using an extensive series of checks performed by the function validate_abc() (Figure 2D). This function tests every component of the ABC pipeline step by step, providing a detailed summary of the validation process and highlighting any potential issues before any computational effort is invested on full-scale simulations.

Having made sure that every step of the inference workflow is in place, users proceed with data simulation (simulate_abc(), Figure 2E) and posterior distribution inference (run_abc(), Figure 2F). Because large-scale ABC workflows are extremely computationally intensive, *demografr* utilizes the functionality of the R package *future* to allow users to parallelize their simulations, simply by instructing the plan() function to define the number of CPUs to use for computation, with no additional work (Figure 2E).

The final product of a *demografr* ABC workflow is a single R object, produced by the run_abc() routine. This object encapsulates information about the joint posterior distribution of parameters inferred from the model. The package provides a range of functions for examining this posterior distribution, both visually and by extracting summary statistics and posterior samples in a tabular form (Figure 2G, H). Furthermore, *demografr* provides an implementation of a generic R method predict() which will, when applied to an abc object, perform another automated batch of simulations for posterior predictive checks which can be visualized using a dedicated function (Figure 2I). Additional diagnostic functions, largely serving as convenient wrappers to methods of the *abc* R package, are available in the *demografr* library.

### Example 2: Parameter grid search

In Figure 3, we demonstrate the features of *demografr* for grid-based parameter sweeps using a practical example from an earlier study led by one of the authors (Petr et al. 2019). Briefly, significant debates have surrounded the putative long-term monotonic decline of Neanderthal ancestry observed in ancient DNA time series data from humans [38] and its incompatibility with models of selection against introgressed DNA investigated at that time [39,40]. To address this issue in the past, Petr et al. utilized large-scale neutral simulations of expected trajectories of introgressed Neanderthal ancestry across various relevant parameters of anatomically modern human (AMH) demographic histories and discovered that the apparent decline in Neanderthal ancestry observed in empirical data is very likely a statistical artifact of gene-flow events between AMH populations following the Neanderthal introgression. The study’s simulation workflow required a combination of shell scripting to orchestrate Python simulations written in *msprime* as well as data visualization code, all tailored for one specific research question at hand (github.com/bodkan/nea-over-time). If a different project were to perform a similar simulation study (different species, different summary statistics of interest, etc.), this would require writing new scripts, largely from scratch. Using *demografr*, one can replace the laborious process involved in the original study by a few dozen straightforward lines of R code, as demonstrated here (Figure 3).

First, we note that most of the workflow of the grid-based approach (Figure 3) is largely the same as the previous example of a toy ABC pipeline (Figure 2). The main difference lies in the specification of model parameters as a data frame with discrete combinations of parameter values (Figure 3C) instead of prior specification in the previous example (Figure 2B) and executing the simulations across a given number of replicates via a simulation routine, simulate_grid() (Figure 3C) instead of simulate_abc() (Figure 2E). The reader will note that the model specification (an R function called model, Figure 3A), a list of summary statistic functions (Figure 3B) and, to a large extent, even the simulation routine (Figure 3C) remain in the same format. Consistency and modularity were important design goals of *demografr*, so as to enable exchangeability of individual model components (and even simulation routines) to investigate various modeling approaches.

The outcome of the parameter sweep simulations is, yet again, a standard R data frame (Figure 3C) in a *tidy* format [41]. The values of summary statistics are stored as individual list-columns of this data frame (each column name matching the name of a summary statistic function given in the input named R list Figure 3B). As such, interactions between model parameters and their influence on the behavior of the summary statistics can be immediately inspected with standard visualization and data analysis functionality of R (Figure 3D, E).

### Example 3: Fully customized ABC workflow

Although *slendr* offers many convenient features for building “standard” population genetic models (specifically, Wright-Fisher neutral demographic models), the range of possible models for fitting evolutionary genetic data is, of course, much wider. In situations in which the *slendr* package might not be sufficient as a modeling framework, researchers could still be interested in using *demografr* for building inference workflows, but utilize their own, custom-designed, Python *msprime* or SLiM scripts as simulation “engines” instead of *slendr*. Our final example, shown in Figure 4, showcases this possibility.

**Figure 4.**
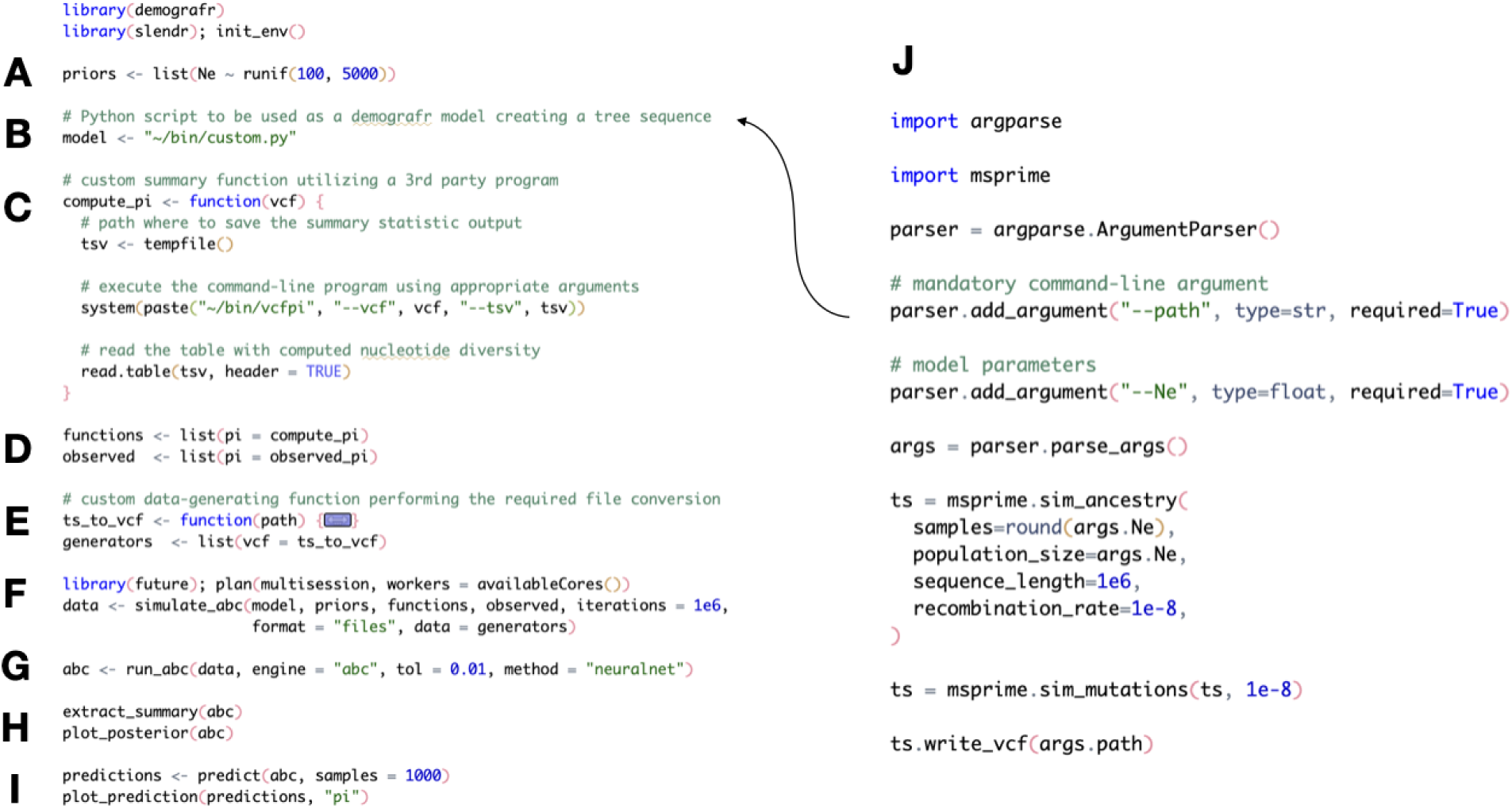
Basic workflow demonstrating individual steps (A-I) of a typical ABC workflow using the *demografr* package which relies on a pure Python simulation script (B) and a third-party summary statistic program (C) for generating simulated data. Complete reproducible code for this example, as well as additional possibilities (such as running SLiM simulations instead) are shown in a corresponding vignette at https://bodkan.net/demografr.

First, users provide a path to a custom Python *msprime* or SLiM script (Figure 4B, J), to be used as an engine which will simulate data given some values of parameters. Unlike the standard *slendr*-based model interface, these parameters (whose priors are specified using the same domain-specific syntax, Figure 4A) are intended to be passed as command-line arguments of this customized script (Figure 4J). Summary statistic functions are specified in exactly the same way as with default *demografr* workflows, in the form of two named R lists (Figure 4D). Beyond being able to use non-*slendr* simulation scripts, users also have the possibility to opt-out of using *slendr*-specific tree-sequence functions entirely, and instead provide their own means to compute summary statistics. In this example, the hypothetical ABC inference workflow relies on a third-party software for computing genetic diversity from a VCF file (a toy command-line utility implemented for this example for demonstrative purposes), bypassing the *slendr*-based summary statistic computation entirely (Figure 4C). However, because *demografr* cannot make any assumptions about the format of the data produced by the custom simulation script or that expected by the statistical third-party software (Figure 4C), users must also provide a “data-generating” function, which is then utilized by *demografr* to connect the two components of an inference pipeline together (Figure 4E). The reader will notice that although both the simulation (Figure 4B) and summary statistic computation (Figure 4C) use fully customized third-party software, the form of the overall workflow remains effectively the same—*demografr* takes all the provided building blocks of the pipeline and orchestrates the execution of the simulations in a fully automated, parallelized manner (Figure 4F). The outcome of the inference is, yet again, an abc object containing the inferred joint posterior distribution (Figure 4G), which can be examined, visualized, and processed (Figure 4H, I) in exactly the same manner as shown in the standard *slendr*-based workflow in the first example (Figure 2).

It is important to note that although we have focused on an inference workflow based on neutral simulation using *msprime*, the flexibility of the *demografr* approach can handle effectively any kind of simulation. Simulation “engines” in the form of SLiM scripts implementing complex non-Wright-Fisher models, selection scenarios, and spatial models, are all supported, as long as they are implemented as command-line scripts which accept model parameters as command-line arguments, and save their results as files in location specified as the --path argument (Figure 4J), so that *demografr* routines can pick them up for downstream inference steps.

## Conclusion

The ever expanding amount of genomic data available for different species has revolutionized our understanding of evolutionary histories across the entire tree of life, making it possible to investigate evolutionary models of increasing complexity. As a result, simulation-based techniques such as Approximate Bayesian Computation (ABC) are becoming even more appealing, owing to their ability to perform parameter inference even in situations in which the complexity of a model makes likelihood approaches impractical or outright impossible [6,8,10,42,43]. However, simulation-based inference is notoriously challenging due to the programming efforts needed to develop simulation and statistical code tailor-made for each specific project. As the opportunity and need to address complex modeling problems increase, it is worth the effort to design novel tools that address both of these problems at once, not least due to frequent reproducibility issues with complex scientific workflows [44].

Our recent development of the *slendr* simulation framework [29] has been intended as a first step towards making population genetic modeling and simulation as straightforward, concise, and reproducible as possible, both for novice researchers and experts alike, joining similar efforts in the broader population genetic community [14,15]. A particular strength of *slendr* is that it requires virtually no prior programming knowledge, beyond the ability to call R functions and basic work with data frames, allowing researchers to focus exclusively on the modeling problem at hand instead of dealing with technicalities. The new package presented in this paper, *demografr*, aims to extend this philosophy even further, towards simulation-based inferences, building on top of modeling components established by the *slendr* package.

Our primary design focus behind *demografr* was to eliminate friction between disparate tools and programming frameworks, offering a streamlined programmable interface to develop simulation-based inference workflows using currently popular techniques such as ABC. Additionally, *demografr* provides individual building blocks allowing the development of entirely new inference routines. In the future, we envision users will be able to swap inference procedures in their *demografr* workflows by changing a single line of code, thus exploring the possibilities and limitations of different approaches to the modeling study at hand with little to no extra coding. Depending on community involvement and feedback, these experimental inference routines could be based on techniques such as Bayesian parameter optimization [31] or random forests [33]. This could also allow a future study which would investigate the tradeoff involved in the choice of various inference techniques depending on the features of a given model under study, the dimensionality of the parameter space, and the choice and characteristics of population genetic summary statistics. Establishing these guidelines will be very important. Although likelihood-free simulation-based approaches such as ABC make it theoretically (and naively) possible to estimate nearly any kind of model regardless of its complexity, extreme care must be taken when performing inference in practice, as the dimensionality of the parameter space can get easily out of hand. Furthermore, even relatively simple models could be ill-posed and potentially non-identifiable and rely on strong assumptions, all of which must be kept in mind when interpreting inferred parameters or deploying models for prediction [36,45,46]. Until now, a thorough exploration of these issues in simulation-based studies has been quite challenging due to the programming efforts required to program even a standard inference pipeline focused on a single inference approach like ABC. The ease of use of *demografr* will significantly decrease the barrier for understanding the limits of various approaches to simulation-based inference.

Despite these pitfalls (which generally apply to statistical modeling in the broadest sense), we believe that *demografr* will not only help the application of simulation-based inference to areas where it has already proven to be a powerful modeling tool [10,43], but will also significantly contribute to approaching problems where simulation-based inference in general has not yet been widely applied, despite its potential benefits. One topic of promising application could be in the area of spatial population genetics. In particular, Identity-by-Descent (IBD) tracts have recently become a powerful source of information about population and social dynamics, especially given the ever-increasing number of available ancient genomes [47].

However, in most such studies, IBD sharing patterns mostly serve a descriptive purpose, and their potential as a powerful summary statistics to elucidate patterns of spatial mobility and social dynamics have yet to be fully explored. In this context, recent development in efficient extraction of IBD tracts from tree-sequence data [48] would be very fruitful, particularly due to *demografr*’s already established tree-sequence interface via the *slendr* package and its possibilities for spatially-explicit simulations. Another promising avenue for leveraging *demografr* in simulation-based inference could be modeling natural selection. Recently, work has begun in *slendr* to implement support for overriding its default neutral mode of operation with fully customizable “SLiM extensions”, allowing users to leverage its powerful interface for easy encoding of complex (neutral) demographic histories while embedding arbitrary SLiM natural selection code within them. Given that *demografr* accepts any kind of *slendr* model as a basis for inference, and given that ABC has been successfully applied to selection in complex demographic histories using traditional software tools [49–51], *demografr* could provide a significant contribution by making these classes of inference workflows easier to program.

Our simulation software framework *slendr* has already shown promise to be a convenient software tool useful not only for unlocking new population genetic insights [52], but also in the classrooms [53]. After all, its features that make it easy for novice researchers and students to get up to speed very quickly, taking them from a spark of a model idea to fully functioning simulation model, are the same reasons why *slendr* helps expert researchers to be more efficient in their daily work. We hope that the *demografr* package will follow in its footsteps, and help to make simulation-based inference more accessible, more powerful, and more reproducible.

## Acknowledgements

M.P. and F.R. were supported by a Novo Nordisk Fonden Data Science Ascending Investigator Award (NNF22OC0076816) and by the European Research Council (ERC) under the European Union’s Horizon Europe program (grant agreement No. 101077592). F.R. was also supported by the ERC under the Horizon 2020 program (grant agreement No. 951385).

## Author contributions

M.P. designed and implemented the software, I.M.P. contributed to testing, F.R. provided advice and support. All authors contributed to writing the manuscript.

